# Map of SARS-CoV-2 spike epitopes not shielded by glycans

**DOI:** 10.1101/2020.07.03.186825

**Authors:** Mateusz Sikora, Sören von Bülow, Florian E. C. Blanc, Michael Gecht, Roberto Covino, Gerhard Hummer

**Affiliations:** Max Planck Institute of Biophysics, Max-von-Laue-Straße 3, 60438 Frankfurt am Main, Germany; Frankfurt Institute for Advanced Studies, Ruth-Moufang-Straße 1, 60438 Frankfurt am Main, Germany; Institute of Biophysics, Goethe University Frankfurt, Max-von-Laue-Straße 1, 60438 Frankfurt am Main, Germany

## Abstract

The severity of the COVID-19 pandemic, caused by the SARS-CoV-2 coronavirus, calls for the urgent development of a vaccine. The primary immunological target is the SARS-CoV-2 spike (S) protein. S is exposed on the viral surface to mediate viral entry into the host cell. To identify possible antibody binding sites not shielded by glycans, we performed multi-microsecond molecular dynamics simulations of a 4.1 million atom system containing a patch of viral membrane with four full-length, fully glycosylated and palmitoylated S proteins. By mapping steric accessibility, structural rigidity, sequence conservation and generic antibody binding signatures, we recover known epitopes on S and reveal promising epitope candidates for vaccine development. We find that the extensive and inherently flexible glycan coat shields a surface area larger than expected from static structures, highlighting the importance of structural dynamics in epitope mapping.

## INTRODUCTION

The ongoing COVID-19 pandemic, caused by the SARS-CoV-2 coronavirus, has emerged as the most challenging global health crisis within a century [1]. As such, the development of vaccines and antiviral drugs effective against SARS-CoV-2 is absolutely urgent. As for other enveloped viruses [2], the primary vaccine target is the trimeric spike (S) protein in the envelope of SARS-CoV-2. S mediates viral entry into the target cell [3–7]. After binding to the human angiotensin-converting enzyme 2 (ACE2) receptor, the ectodomain of S undergoes a drastic transition from a prefusion to a postfusion conformation. This transition drives the fusion between viral and host membranes, which triggers internalization of SARS-CoV-2 via endocytic and possibly non-endocytic pathways [8, 9]. Locking the conformation of S or blocking its interaction with ACE2 would prevent cell entry and infection. However, the dense glycan coat of S effectively shields the virus from an immune response and hinders pharmacological targeting.

A detailed understanding of the exposed viral surface is instrumental in vaccine design [10]. Thanks to the extraordinary response of the global scientific community, we already have atomistic structures of S [6, 7, 11, 12] and detailed views of the viral envelope [13–16]. However, static structures do not capture conformational changes of S or the motion of the highly dynamic glycans covering it. Molecular dynamics (MD) simulations add a dynamic picture of S and its glycan protective shield [17–19]. Intriguingly, Amaro’s group predicted that glycans also play a crucial role in the infection mechanism [18]. Recent experiments validated these results [20], confirming the potential of accurate atomistic models.

Here, we report on the ~2 μs-long MD simulation of a full-length atomistic model of four S trimers in the pre-fusion conformation, giving us in aggregate ~8 μs of S dynamics. The model includes the transmembrane do main (TMD) embedded in a realistic membrane, along with realistic post translational modification patterns, i.e., glycosylation of the ectodomain and palmitoylation of the TMD. Although independently developed, our S protein model and its structural dynamics are in quantitative agreement with recent high-resolution electron cryotomography (cryoET) images [15].

We identify possible immunogenic epitopes on SARS-CoV-2 S by combining information on steric accessibility and structural flexibility with bioinformatic assessments of sequence conservation and epitope characteristics. We recover known epitopes in the ACE2 receptor-binding domain (RBD) and identify several epitope candidates on the spike surface that are exposed, structured, and conserved in sequence. In particular, target sites for antibodies emerge in the functionally important S2 domain harboring the fusion machinery. We propose the structural domains presenting these epitopes as possible immunogens.

## METHODS

### Full-length molecular model of SARS-CoV-2 S glycoprotein

Our simulation system contained four membrane-embedded SARS-CoV-2 S proteins assembled from resolved structures where available and models for the missing parts (SI Appendix, Fig. S5). The spike head was modeled based on a recently determined structure (PDB ID: 6VSB[6]) with one RBD domain in an open conformation and glycans modeled according to [21]. The stalk connecting the S head to the membrane was modeled *de novo* as trimeric coiled coils, consistent with an experimental structure of the HR2 domain in SARS-CoV S (PDB ID: 2FXP[22]). The TMD as well as the cytosolic domain were modeled *de novo*. See SI Appendix, supplementary methods for further details and SI Appendix, Fig. S6 for a view of the final model.

### Molecular dynamics simulations

We assembled four membrane-embedded full-length S proteins to form one large membrane patch with proteins spaced at about 15 nm distance [23, 24], totaling ~4.1 million atoms in the system. We performed MD simulations of the four S proteins for 1.93 μs in the *NpT* ensemble with GROMACS 2019.6 [25]. We used the CHARMM36m protein and glycan force fields [26–28], in combination with the TIP3P water model, and sodium and chloride ions (150 mM).

### Rigidity analysis

We quantified the local rigidity in terms of RMSF values. For each frame, the C_α_ atoms of residues within 15 Å of the residue of interest were rigid body aligned to the average structure. The RMSF values were then averaged over the C_α_ atoms, weighted by the relative surface area of each residue [29]. These flexibility profiles were averaged over the four spike copies and three chains. The local rigidity was then defined as the reciprocal of the flexibility.

### Accessibility analysis

The accessibility of the S protein surfaces was probed by illuminating the simulation protein in diffuse light, as detailed below, and by rigid body docking of the Fab of the antibody CR3022 [30], as detailed in the SI Appendix. For the illumination analysis, rays of random orientation emanate from a half-sphere with radius 25 nm around the center of mass of the protein. They are absorbed by the first heavy atom they pass within 1.5 Å

Simulation structures collected at 10 ns intervals were each probed with 10^6^ rays. To quantify the effect of glycosylation, the analysis was performed with and without including the glycan shield. The fraction of rays absorbed was used as one measure of accessibility, and possible contact with an Fab (SI Appendix) as the other.

### Sequence variability analysis

To estimate the evolutionary variability of the S protein, we analyzed the aligned amino acid sequences released by the GISAID initiative on 25 May 2020 (https://www.gisaid.org/). We first built the consensus sequence with the most common amino acid (the mode) at each position across the whole data set. We then kept only 1273 amino acid long sequences, and filtered out corrupted sequences by discarding those having a Hamming distance from the cosnensus larger than 0.2. With the remaining 30,426 sequences, we estimated the conservation at each position [31]. Our conservation score is defined as the normalized difference between the maximum possible entropy and the entropy of the observed amino acid distribution at a given position, cons (*i*) = 1 + Σ_*k*_ *p*_*k*_ (*i*) log *p*_*k*_ (*i*) */* log 20, where *p*_*k*_ (*i*) is the probability of observing amino acid *k* at position *i* in the sequence.

### Sequence-based epitope predictions

We estimated the epitope probability prediction by using the BepiPred 2.0 webserver (http://www.cbs.dtu.dk/services/BepiPred/), with an Epitope Threshold of 0.5 [32]. BepiPred 2.0 uses a random forest model trained on known epitope-antibody complexes.

### Consensus score for epitope prediction

We integrated the information of the different analyses into the consensus epitope score. We first applied a 3D Gaussian filter with *σ* = 5 Å to the ray and docking scores. We then mapped each score to the interval [0, 1], with outliers mapped to the extremes listed in SI Appendix, Table S1. Finally, we multiplied the individual scores together to obtain the consensus score, which was also mapped to [0, 1].

## RESULTS

### Model of full-length S

As basis for our search for possible epitopes, we constructed a detailed structural model of glycosylated full-length S. Whereas high-resolution structures of the S head are available [6, 7], the stalk and membrane anchor have so far not been resolved at atomic level. Moreover, the glycosylation partially resolved in the S head structures may differ from that under infection conditions because of its passage through an intact Golgi in the expression system [15].

We built a model of the complete S by combining experimental structural data and bioinformatic predictions. Our full-length model of the S trimer consists of the large ectodomain (residues 1-1137) forming the head, two coiled-coil domains, denoted CC1 (residues 1138-1158) and HR2 (residues 1167-1204), forming the stalk, the *α*-helical TMD (residues 1212-1237) with flanking amphipathic helices (1243-1255) and multiple palmitoylated cysteines, and a short C-terminal domain (residues 1256-1273). The model fits high-resolution cryoET electron density data of S proteins on the surface of virions extracted from a culture of infected cells remarkably well [15].

### Multi-microsecond atomistic MD simulations reveal dynamics of S and its glycan shield

We performed a ~2 μs long atomistic MD simulation of a viral membrane patch with four flexible S proteins, embedded at a distance of about 15 nm [23, 24] (Fig. 1). During the simulation, the four S proteins remained folded and anchored in the membrane with well separated TMDs. The S heads tilted dynamically and interacted with their neighbors (SI Appendix, Movie S1). High-resolution cryoET images [15] and a recent MD study [18] independently revealed significant head tilting associated with flexing of the joints in the stalk, in strong support of our observations. Being highly mobile, the glycans on the surface of S cover most of its surface (Fig. 2*A-C*).

**FIG. 1.**
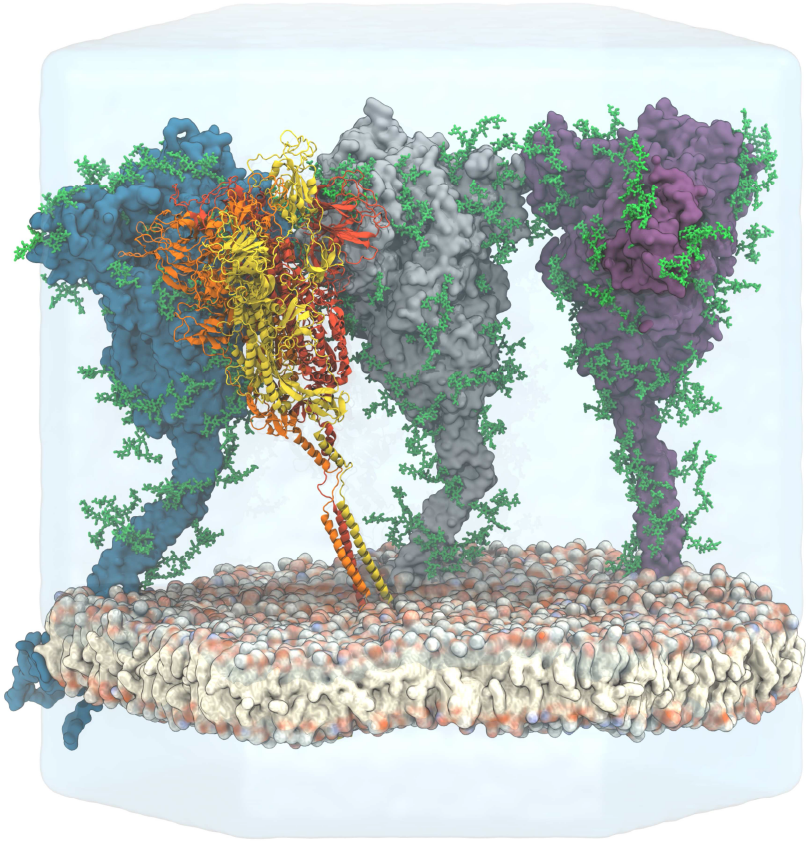
View of the simulated atomistic model containing four glycosylated and membrane-anchored S proteins in a hexagonal simulation box. Three proteins are shown in surface representation with glycans represented as green sticks. One protein is shown in cartoon representation, with the three chains colored individually and glycans omitted for clarity. Water (transparent) is only shown for the lower and back half of the hexagonal box, and ions are omitted for clarity.

**FIG. 2.**
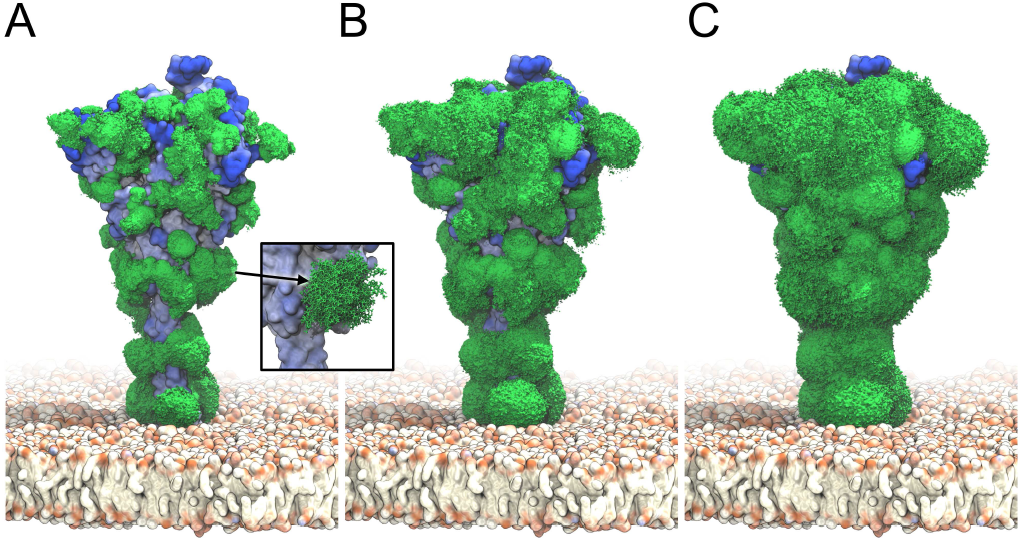
S glycan dynamics from ~2 μs MD simulations. Time-averaged glycan electron density isosurfaces are shown at high (*A*), medium (*B*), and low (*C*) contour levels, respectively. The blue-to-white protein surface indicates high-to-low accessibility in ray analysis. (Inset) Snapshots (sticks) of a biantennary, core-fucosylated and sialylated glycan at position 1098 along the MD trajectory.

### Antibody binding sites predicted from accessibility, rigidity, sequence conservation, and sequence signature

#### Accessibility of the S ectodomain

Antibody binding requires at least transient access to epitopes. The general accessibility of S on the viral membrane and the surface coverage by glycans was extracted from the MD simulations by (1) ray and (2) antigen-binding fragment (Fab) docking analyses. In the ray analysis, we illuminated the protein model by diffuse light; in the Fab docking analysis, we performed rigid body Monte Carlo simulations of S and the SARS-CoV-2 antibody CR3022 Fab to determine the steric accessibility to an antibody Fab. To account for protein and glycan mobility, we performed both analyses individually for 4 × 193 snapshots taken at 10 ns time intervals from the 1.93 μs MD simulation with four glycosylated S proteins.

The glycan shield dramatically reduces the accessibility to the S surface (Fig. 3*A,B*). Ray accessibility time traces show that the accessibility of a given site varies considerably in time (SI Appendix, Fig. S1). Even though glycans cover only a small fraction of the protein surface at a given moment (Fig. 1), their high mobility leads to a strong effective shielding of S (Fig. 2). A comparison of the Fab docking results for glycosylated and unglycosylated S further illustrates this effect (SI Appendix, Fig. S2). Ray and docking analyses show that glycans cause a reduction in accessibility by about 35% and 80%, respectively. The most marked effect occurs in the HR2 coiled coil close to the membrane. Without glycosylation, HR2 is fully accessible; with glycosylation, HR2 becomes inaccessible to Fab docking (Fig. 4*A,B*). Whereas small molecules may interact with the HR2 protein stalk, antibodies are blocked from surface access, in agreement with recent simulations by Casalino et al. [18].

#### Rigidity of S

Structured epitopes are expected to bind strongly and specifically to antibodies. By contrast, mobile regions tend to become structured in the bound state, entailing a loss in entropy and may not retain their structure when presented in a vaccine construct. With the aim of eliciting a robust immune response, we chose to include rigidity in our epitope score. Here, we are less concerned with the large-scale conformational dynamics associated with the flexible hinges in the stalk and membrane anchor, as analyzed in another paper [15]. Instead, we concentrate on motions of domains on the scale of about 1 nm. For this, we determined the root-mean-square fluctuations (RMSF) locally by superimposing local protein regions and converting the RMSF into a rigidity score, as described in Methods.

**FIG. 3.**
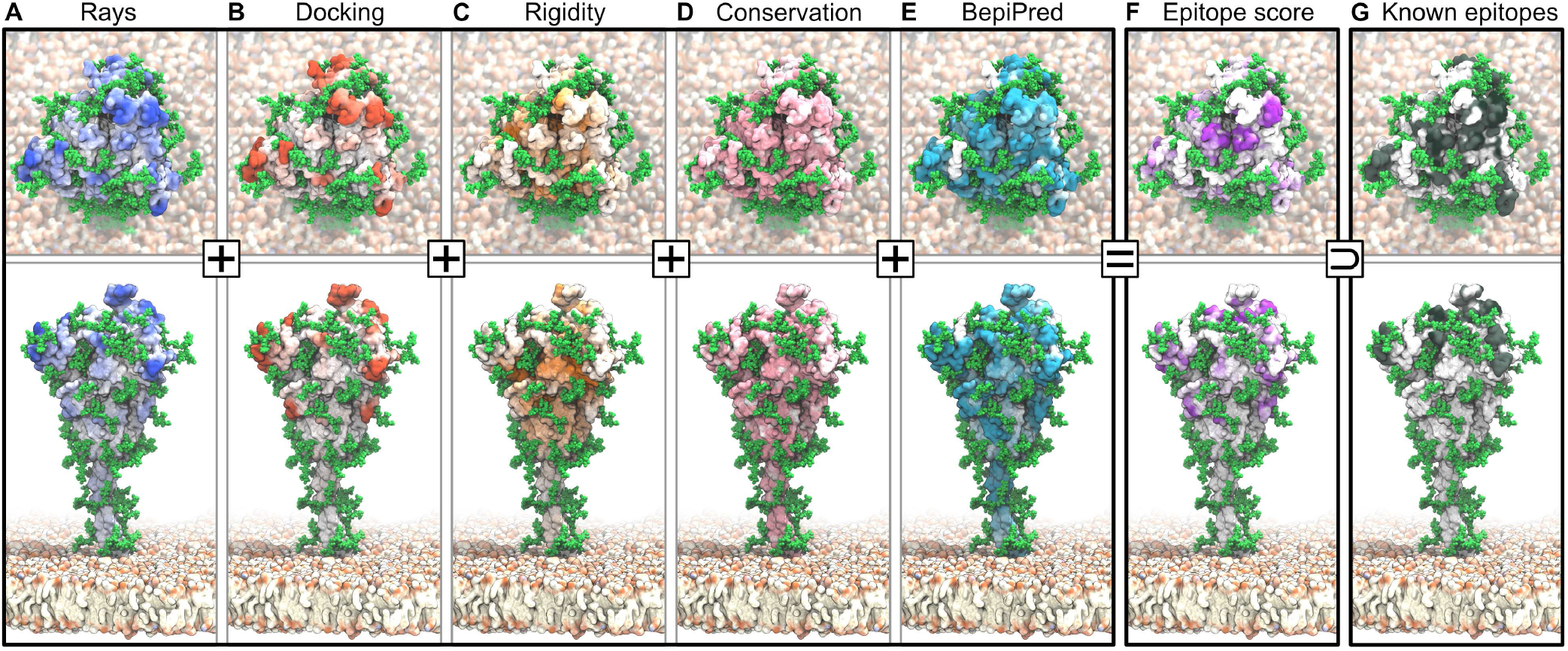
Epitopes identified from MD simulations and bioinformatics analyses. Accessibility scores from (*A*) ray analysis and (*B*) Fab rigid body docking are combined with (*C*) rigidity scores, all averaged over 4 × 1.93 μs of S protein MD simulations. Also included are (*D*) a sequence conservation score [31], and (*E*) BepiPred-2.0 epitope sequence-signature prediction. (*F*) Combined epitope score. (*G*) Binding sites of known neutralizing antibodies. Higher color intensity in *A-F* indicates a higher score and higher color intensity in *G* indicates sites binding to multiple different antibodies.

**FIG. 4.**
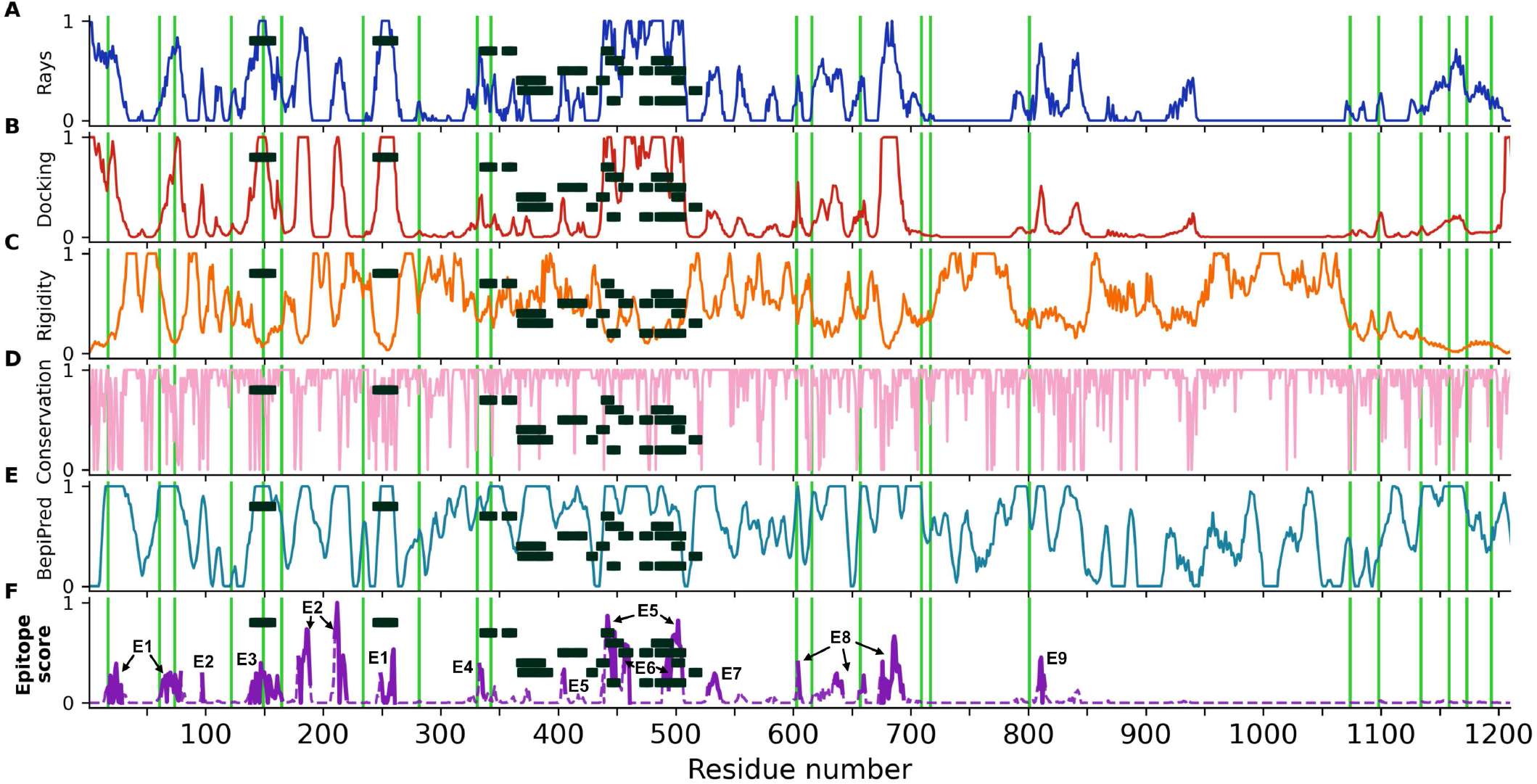
Epitope scores of S ectodomain. Panels (A-F) and colors as in Fig. 3. All values are filtered and normalized (see Methods). Labels E1-E9 in (F) highlight candidate epitopes. Green lines indicate glycosylation sites, and black rectangles known antibody binding sites.

The surface of S presents both dynamic and rigid regions (Fig. 4*C*). Interestingly, the RBD and its surrounding are comparably flexible, consistent with the experimental finding of large differences in the structure of the three peptide chains in open and closed states [7]. By contrast, the protein surface of the S2 domain covering the fusion machinery is relatively rigid (Fig. 4*C*), possibly to safeguard this functionally critical domain in the metastable prefusion conformation.

#### Sequence conservation

Targeting epitopes whose sequences are highly conserved will ensure efficacy across strains and prevent the virus from escaping immune pressure through mutations with minimal fitness penalty. We estimated the sequence conservation from the naturally occurring variations at each amino acid position in the sequences collected and curated by the GISAID initiative (https://www.gisaid.org/). The analysis of 30,426 amino acid sequences revealed that S is highly conserved, with no mutation recorded for 52% of the amino acid positions. As conservation score, we mapped the entropy at each position to the interval between zero and one (see Methods). Even surface regions are mostly well conserved in sequence (Fig. 4*D*).

#### Sequence-based immunogenicity predictor

Conserved, rigid, and accessible regions present good candidates for binding of protein partners in general. To complement this information, we assessed the immunogenic potential based on sequence signatures targeted by antibodies. The epitope like motifs in the S sequence identified using the BepiPred 2.0 server [32] lie scattered across the S ectodomain and include known epitopes (Fig. 3*E* and Fig. 4*E*), but also contain buried regions inaccessible to antibodies.

#### Consensus epitope score

We combined our accessibility, rigidity, conservation, and immunogenicity scores into a single consensus epitope score (Figs. 3*F* and 4*F*). By taking the product of all individual scores, we ensured that epitope candidates have high scores in all features. This rigorous requirement eliminates many candidate sites, mostly because accessibility scores (Fig. 4*A,B*) and the rigidity score (Fig. 4*C*) show opposite trends, in line with the extensive occurrence of flexible loops on the S surface.

Using our consensus score, we identified nine epitope candidates (E1-E9; Fig. 5 and Table I). Epitope candidates E3-E6 recover known epitopes (Fig. 3*F,G* and Fig. 4*F*), achieving residue level accuracy in some cases (SI Appendix, Fig. S3); in addition, we identify epitope candidates E1, E2, and E7-E9. All epitope candidates reside in the structured head of S. By contrast, low accessibility and high flexibility in the hinges [15] give the stalk low overall epitope scores.

**FIG. 5.**
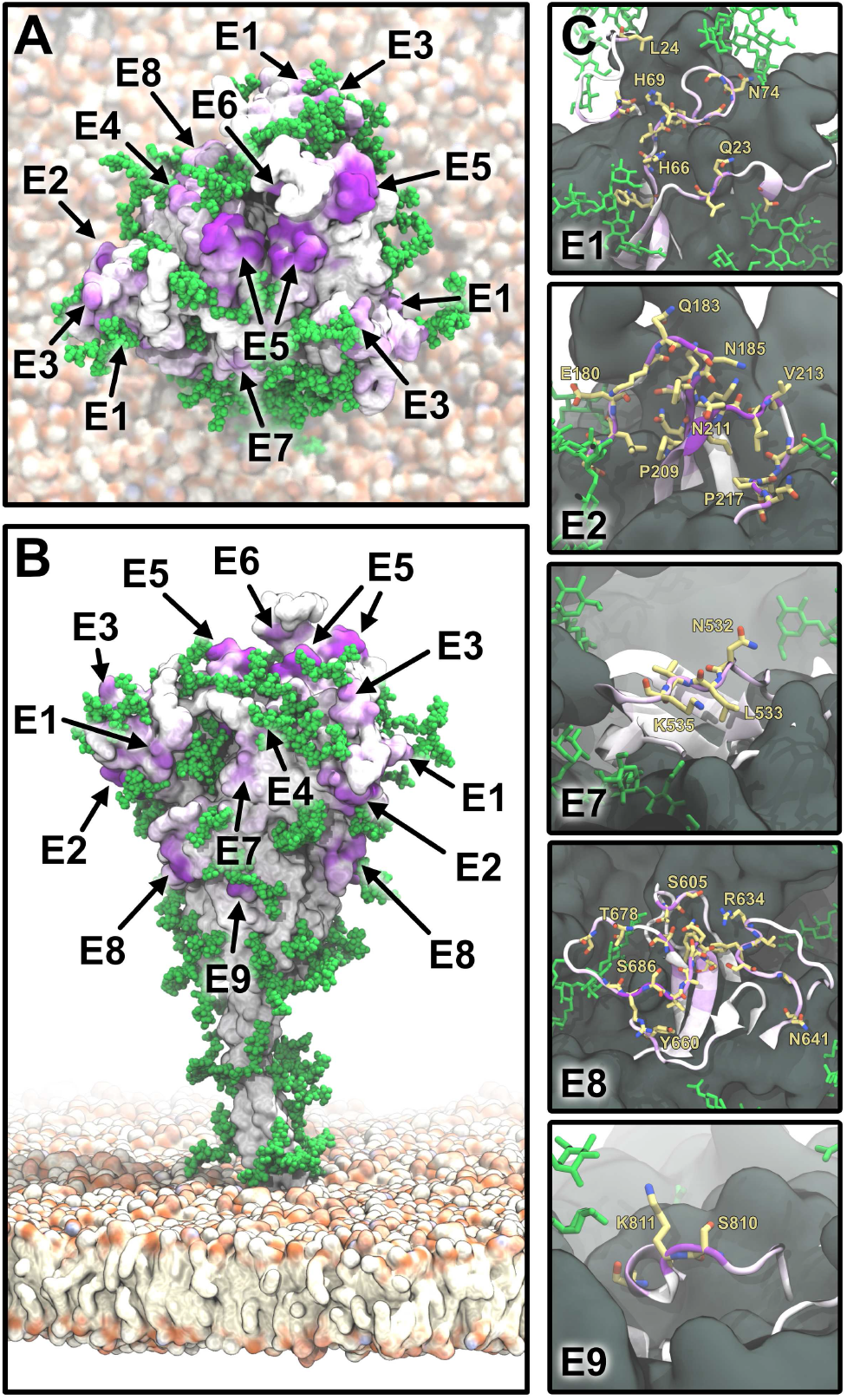
S epitopes. (*A*) Top view of S represented as in Fig. 3*F*. Epitope candidates are labeled according to Table I. (*B*) Side view with coloring and labels as in A. (*C*) Zoom-ins on epitope candidates (E1, E2, E7-E9) in a cartoon representation and colored according as in A. Residues with an epitope consensus score >0.2 are shown in yellow licorice representation.

**TABLE I.** Epitopes shown in Figs. 3–5.

| Epitope | Residues |
| --- | --- |
| E1 | 15-28, 61-79, 248-261 |
| E2 | 96-97, 178-188, 209-219 |
| E3 | 137-164 |
| E4 | 332-346 |
| E5 | 403-405, 438, 440-450, 494-505 |
| E6 | 455-460, 489-493 |
| E7 | 527-537 |
| E8 | 603-605, 634-641, 656-660, 674-693 |
| E9 | 808-813 |

### DISCUSSION

#### Validation through identification of known epitopes

Even though SARS-CoV-2 was identified only a few months ago, several groups have already reported on antibodies binding the SARS-CoV-2 S protein [30, 33–38]. Most notably, Yuan and co-workers structurally characterized the binding of SARS-CoV-neutralizing antibody CR3022 to the SARS-CoV-2 S protein ectodomain [30, 33, 39]. Their structure reveals an epitope distal to the ACE2 binding site that requires at least two of the S protomers to be in the open conformation to permit binding without steric clash. Interestingly, while our simulations do not probe the doubly open configuration, the epitope reported by Yuan et al. [30] is still successfully identified with a significant consensus score. Moreover, epitopes for other reported antibodies H104 [34], CB6 [35], P2B-2F6 [36], S309 [38], and 4A8 [40] also match regions of high consensus score. In particular, our candidate epitopes E5 and E6 overlap with the reported binding sites in the RBD for neutralizing antibodies. We conclude that our epitope identification methodology is robust.

#### Dependence on detailed glycosylation pattern

The complexity and variability of S glycosylation *in situ* remains poorly understood. Mass spectrometry on recombinant S confirmed its extensive glycosylation [21]. CyoET images of intact viral particles revealed branching points in the glycans [15], indicative of complex glycans. We addressed this uncertainty by repeating our docking accessibility analysis for different glycosylation patterns. We considered pruned (mannose-only) glycans by removing fucose, sialic acid, and galactose. Remarkably, this pruned glycan shield impedes Fab accessibility almost as effectively (~60%) as the full shield (~80%), even if epitopes E8 and E9 become somewhat more exposed with trimmed glycans (SI Appendix, Fig. S2). Overall, we conclude that even a light glycan coverage strongly reduces the antibody accessibility of the protein.

#### Structural and dynamic characteristics of candidate epitopes

Epitopes E1-E3 are part of the N-terminal domain, which is formed mostly of antiparallel *β* sheets (residues 1-291). All three epitopes include flexible loops and folded *β* strands (SI Appendix, Fig. S4*A*). Epitope E4 is located on a two-turn *α*-helix flanked by a short twin *α*-helix and lying on a five-strand antiparallel *β*-sheet. This arrangement provides the epitope with remarkable stability (SI Appendix, Fig. S4*B*). Epitopes E5 and E6 are located on the apical part of S in the RBD, and are composed mostly of flexible loops. E5 and E6 jointly span a contiguous surface in chain A, which is in the open conformation. By contrast, in the closed chains B and C, this surface is altered and E6 is buried (SI Appendix, Fig. S4*C*). The epitope E7 is part of a stable helix that connects neighboring *β*-sheets (SI Appendix, Fig. S4*D*). E8 comprises two quite long and flexible loops (residues 634-641 and 674-693), and two shorter and less flexible ones (SI Appendix, Fig. S4*E*). Finally, E9 is located on a short and flexible loop (SI Appendix, Fig. S4*F*).

#### Guidance for immunogen and vaccine design

Having identified accessible, relatively structured, and conserved epitope sequences, one of the challenges is to present these epitopes in an immunogenic manner to induce a robust antibody response. The structure of SARS-CoV-2 S comprises distinct domains with residue numbers (A) 26-290, (B) 591-699, and (C) 1071-1138. Domains A and B encompass the predicted epitopes E1, E2 and E3, respectively, and E8 is contained within domain B. We speculate that these domains may fold independently, possibly after suitable sequence redesigns, and could thus be used to present the epitopes faithfully.

#### Glycans as epitopes

There have been reports of glycan mediated antibody binding to SARS-CoV-2 S [18] and to HIV-1 Env [41]. While this could open up possibilities for epitope binding, the natural variability of the glycan shield [21], along with its extensive structural dynamics demonstrated here, currently preclude a systematic search for glycan-involving epitopes. Moreover, with human and viral proteins carrying chemically equivalent glycan coats, the risk of autoreaction is significant [41]. Therefore, we concentrated here on amino acid epitopes.

## CONCLUSIONS

We identified epitope candidates on the SARS-CoV-2 S protein surface by combining accurate atomistic modelling, multi-microsecond MD simulations, and a range of bioinformatics and analysis methods. We concentrated on sites that are accessible to antibodies, unencumbered be the glycan shield, and fairly rigid. We also required these sites to be conserved in sequence and to display signatures expected to elicit an immune response. From all these features, we determined a combined consensus epitope score that enabled us to predict nine distinct epitope sites. Validating our methodology, we recovered five epitopes that overlap with experimentally characterized epitopes, including a “cryptic” site [30].

Highly dynamic glycans shield a large fraction of the S surface. Even though the instantaneous surface coverage of the glycans is low, the long-time average density of few well placed glycans covers most of the protein surface. In particular, only three N-glycosylation sites per protein chain suffice to shield the stalk domain and block antibody binding to this functionally critical part of the protein. New and conflicting reports emerge on the glycan types on the S surface [21, 42], with glycan composition possibly varying from host to host. We considered both light and heavy glycan coverages in our analysis, which should encompass most of the glycan variability. Both extremes show that glycosylation strongly protects S from interactions with antibodies.

The different epitopes we predicted are the starting point to engineering stable immunogenic constructs that robustly elicit the production of antibodies. A fragment-based epitope presentation avoids the many challenges of working with full-length S, a multimeric and highly dynamic membrane protein, whose prefusion structure is likely metastable. Epitopes E1, E2, E3, and E8 are particularly promising candidates. They are located on distinct S domains that could fold independently and present these epitopes in a native like manner. Additionally, epitopes that are distributed on the surface of S will make the onset of resistance due to mutations less likely. The approach we introduced in this paper could be extended to predict epitopes from an integrated analysis of diverse betacoronaviruses, with the ultimate aim of producing a universal vaccine that guarantees broad protection against the whole virus family.

## ACKNOWLEDGEMENTS

We thank Martin Beck, Beata Turoňová, and Philipp Schmalhorst for stimulating discussions, the Max Planck Society for generous support, the Max Planck Computing and Data Facility for providing computational resources, and the Leibniz Supercomputing Centre Munich for the SUPERspike computing allocation. R.C. acknowledges the support by the Frankfurt Institute for Advanced Studies. M.S. acknowledges support by the Austrian Science Fund FWF (Schroedinger Fellowship, J4332-B28). S.v.B. and G.H. acknowledge support by the Human Frontier Science program (RGP0026/2017). M.G. and G.H. acknowledge support by the Landes-Offensive zur Entwicklung Wissenschaftlich-ökonomischer Exzellenz (LOEWE) Dy-naMem program of the state of Hesse.

## Supporting Information

## SUPPLEMENTARY METHODS

### Full-length molecular model of SARS-CoV-2 S glycoprotein

The modeling procedure of the full-length SARS-CoV-2 S glycoprotein is outlined in Fig. S5. We based our model of the SARS-CoV-2 S1/S2 S domain on a recently determined structure (PDB ID: 6VSB[1]). We added missing loops using MODELLER [2]. We modeled the stalk connecting the S head to the membrane as two distinct coiled coils (CCs, henceforth denoted CC1 and HR2) based on CC predictions [3, 4]. CC1 and HR2 at positions 1138-1158 and 1167-1204 are predicted with low and high confidence, respectively. However, since the N-terminal ends of the three helices in CC1 have been resolved in the experimental structures [1, 5], we modeled both segments as trimeric CCs with CCBuilder [6], using the heptad repeat register prediction of [3] and generously extending all termini by several residues to prevent destabilization of the CCs from solvation effects at the termini. Thus, the first model of CC1 comprised residues 1137-1163, while HR2 comprised residues 1161-1214. We then performed 1 μs-long MD simulations of the solvated CC1 and HR2 models individually with procedures and parameter settings as described below. In CC1 and HR2, residues 1138-1158 and 1167-1204 retained stable CC structures, respectively (Fig. S7). The CC structures of snapshots at 390 ns (CC1) and 166 ns (HR2) were integrated into a model of full-length SARS-CoV-2 S.

### Glycosylation of S ectodomain and connector domain

There are 22 N-glycosylation sequons present on the surface of S, all of which have been confirmed recently by mass spectrometry of a recombinant protein [7]. Distinct glycan types are preferred on various sequons, with the majority being oligomannose, followed by sialylated and fucosylated complex glycans and a minority of the hybrid type. Here we selected the most abundant species at each site, as shown in Fig. S8. All fucose residues were linked in α-1,3 position and sialic acid in α-2,3. Consistent with the low glycan occupancy in the structure *in situ* [8], O-glycosylation in positions 323 and 325 was not included. Contrary to some observations [9], the complete glycosylation pattern including heavy glycosylation of the stalk seems to reflect better the situation *in situ* [8].

### Modeling of the transmembrane domain

Lacking a structure for the S transmembrane domain (TMD), we used a hierarchical procedure to model the TMD trimer. Secondary structure predictions revealed that the TMD is likely to be formed of two helical stretches with a long transmembrane helix (residues 1212-1237), followed by a shorter C-terminal helix (residues 1242-1249) with features of an amphipathic helix. The remaining 24 C-terminal residues were predicted as disordered. We hypothesized that the C-terminal helix extends to K1255 and encompasses all cysteine residues, leaving a total of 18 disordered residues at the C-terminus.

We used a manually curated sequence alignment to build a homology model of the S protein TMD trimer helical core (residues 1208-1237) with MODELLER [2]. We palmitoylated all cysteines, inserted the trimer into a lipid bilayer (see below and Table S2), and relaxed the system using molecular dynamics (MD; see parameters below) for 1 μs, to properly equilibrate the relative orientation of the protomers.

Separately, we built an L-shaped TMD monomer model by appending the C-terminal helix (residues 1243-1265, modeled as an ideal α-helix) to the TMD core helix (residues 1208-1237). The C-terminal helix was oriented such that all cysteines pointed into the hydrophobic core of the membrane. The five residues connecting the TMD and C-terminal helix, as well as the 18 C-terminal residues were modeled as unstructured loops using MODELLER [2]. All cysteines were palmitoylated and the monomer was inserted into a lipid bilayer, then relaxed by molecular dynamics for 1 μs for proper positioning of the C-terminal helix with respect to the lipid head groups.

Finally, a TMD trimer model was obtained by structurally fitting the relaxed L-shaped monomer onto the relaxed transmembrane trimer. In two out of three monomers, this resulted in an outward-pointing, clash-free C-terminal helix. In the third monomer, the C-terminal helix was manually rotated around the z-axis to relieve clashes.

### Assembly of full-length S model

A full-length model of S was built by manually matching the separate structural domains using PyMOL [10], and then building missing connecting residues as unstructured linkers with MODELLER [2].

### Membrane lipid composition

Coronaviruses like MERS-CoV and SARS-CoV are assembled in the endoplasmic reticulum (ER) [11]. We therefore modeled the viral envelope with an ER-like composition [12] as detailed in Table S2. The transmembrane domain structures described above were inserted into the ER-like membrane using CHARMM-GUI [13–17].

### Molecular dynamics simulations

Molecular dynamics simulations were performed with GRO-MACS 2019.6 [18], using the CHARMM36m protein and glycan force field [19–21], in combination with the TIP3P water model [22]. Ions parameters were those by Luo and Roux [23].

After energy minimization using the steepest descent algorithm for 55 000 steps, the system was equilibrated in the *NVT* ensemble for 375 ps with a time step of 1 fs, followed by 1500 ps with a time step of 2 fs. In the equilibration runs, the Berendsen thermostat [24] was used for temperature coupling, with the coupling constant τ = 1 ps. After 250 ps, we used the Parrinello-Rahman barostat [25] to apply semiisotropic pressure coupling, using τ = 5 ps and compressibility 4.5 × 10^−5^ bar^−1^. LINCS constraints[26] were applied to all bonds involving hydrogen atoms, allowing us to use a 2 fs integration timestep for equilibration. During equilibration, restraints on positions and dihedrals were gradually decreased from 1000 kJ mol^−1^ nm^−2^ to 0.

Due to the large system size, we adopted specific strategies to enhance the simulation speed during production. We used an integration timestep of 4 fs. All hydrogen masses were doubled, corresponding to deuterium, to avoid instabilities from high frequency vibrations. Cutoffs for non-bonded interactions were set to 1 nm. In addition, temperature control was switched to the V-rescale thermostat [27]. We used MDBenchmark to perform scaling studies and determine the optimal hardware configuration and run settings (MPI ranks/OpenMP threads) [28].

### Rigid-body docking

We probed the steric accessibility for antibody binding using rigid-body docking. The Fab of antibody CR3022 (PDB ID: 6W41 [29]) was used for a coarse-grained rigid-body Monte Carlo (MC) docking analysis, following the procedure described in [30]. Backbone C_α_ atoms were recorded every 10 ns of the MD simulation. Each snapshot was centered in a 24.5 nm × nm × 36 nm orthorhombic simulation box. The Fab was subjected to 2 × 10^5^ translation and rotation MC moves, recorded every 20 moves.

In a first step, we probed the steric accessibility of the protein surface without glycans using rigid-body MC simulations at high temperature (T = 10 000 K). Contacts between the complementarity-determining region of the Fab (heavy chain residues 31-35, 50-65, 95-102; and light chain residues 24-34, 50-56, 89-97) and S were then counted based on a distance criterion of twice the sum of van der Waals (vdW) radii of the amino acids involved in the contact (with radius definitions following [30]). In a second step, we assessed the influence of glycans on the steric surface coverage by excluding all snapshots in which the Fab clashed with glycans. The full glycans and a mannose-only (“pruned”) version of the glycans were considered. Every sugar residue of a glycan was represented by a pseudoparticle positioned at the residue center of mass. The effective vdW radius of this sugar bead was estimated from the sugar residue radius of gyration and found to be roughly equal to the vdW radius of an alanine residue, as defined in [30]. A distance cutoff of the sum of Fab residue vdW radius and glycan (≈ alanine) vdW radius was used to determine clashes.

## SUPPLEMENTARY RESULTS

### Evaluation of the accessibility reduction due to the glycan shield

We quantified the glycan coverage by comparing global accessibility (to rays or to the rigid, coarse-grained Fab) of the S surface with full glycans and without glycans. First, the global accessibility was computed as the sum over all residues of the numbers of hits for a given probing method and glycosylation pattern. Then, we considered the ratio of global accessibility with glycans over global accessibility without glycans. Finally, the relative accessibility reduction due to glycan coverage was taken as the complementary of this global accessibility ratio (relative accessibility reduction = 1 − global accessibility ratio).

## SUPPLEMENTARY TABLES AND FIGURES

**TABLE S1.**
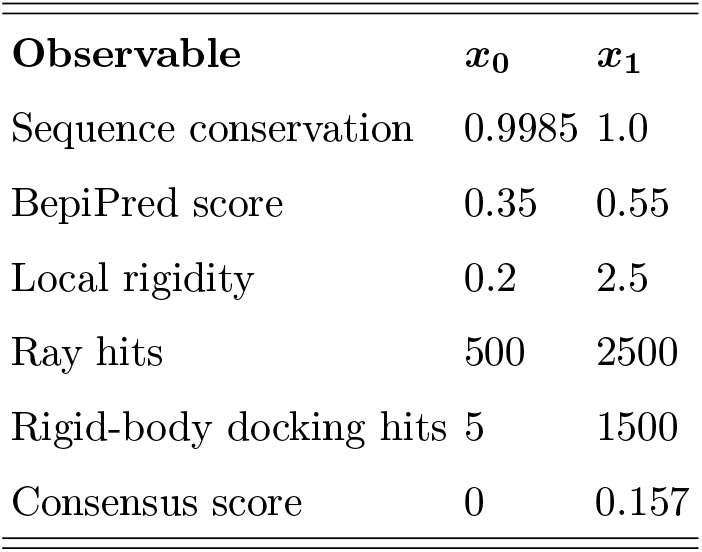
Parameters for mapping of an individual score *x* to the interval [0, 1], with *x* ≤ *x*_0_ mapped to 0, *x* ≥ *x*_1_ mapped to 1, and linear interpolation in between. Numbers are given in the respective units of the corresponding observable.

**TABLE S2.**
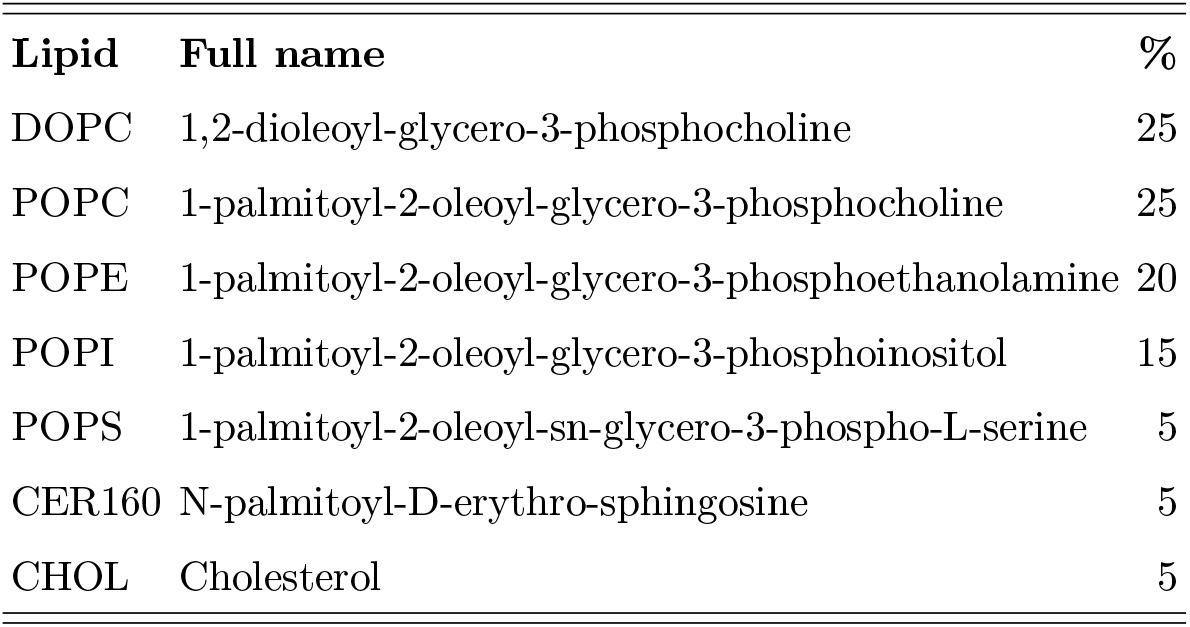
ER-like membrane composition used in the present study

**FIG. S1.**
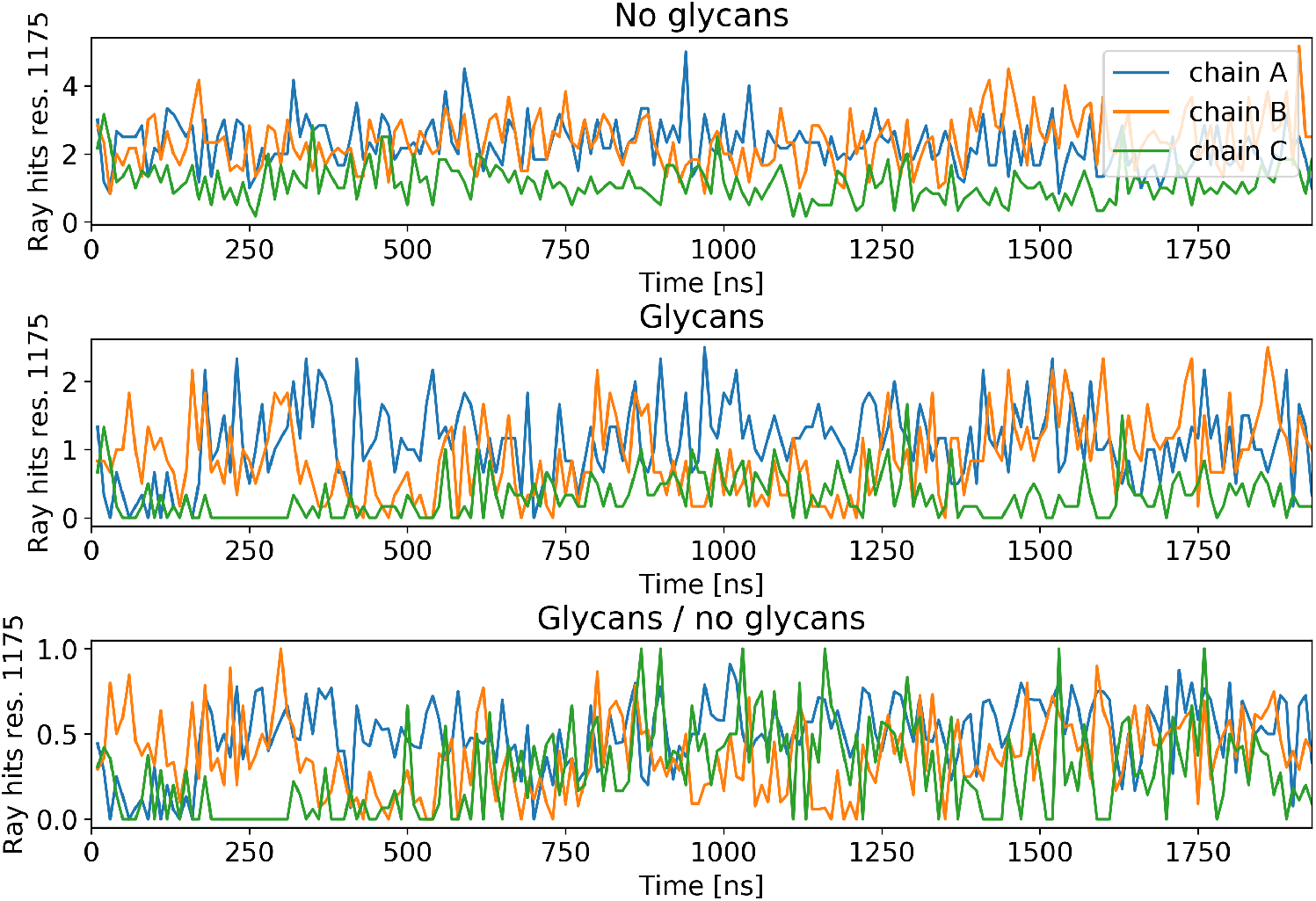
Ray analysis accessibility time traces of residue 1175 in the HR2 domain of S. Mean number of ray hits per timestep. No glycan: Ray analysis without considering glycan shielding. Glycan: Ray analysis taking into account glycan shielding. Glycans / no glycans: Ratio of the above.

**FIG. S2.**
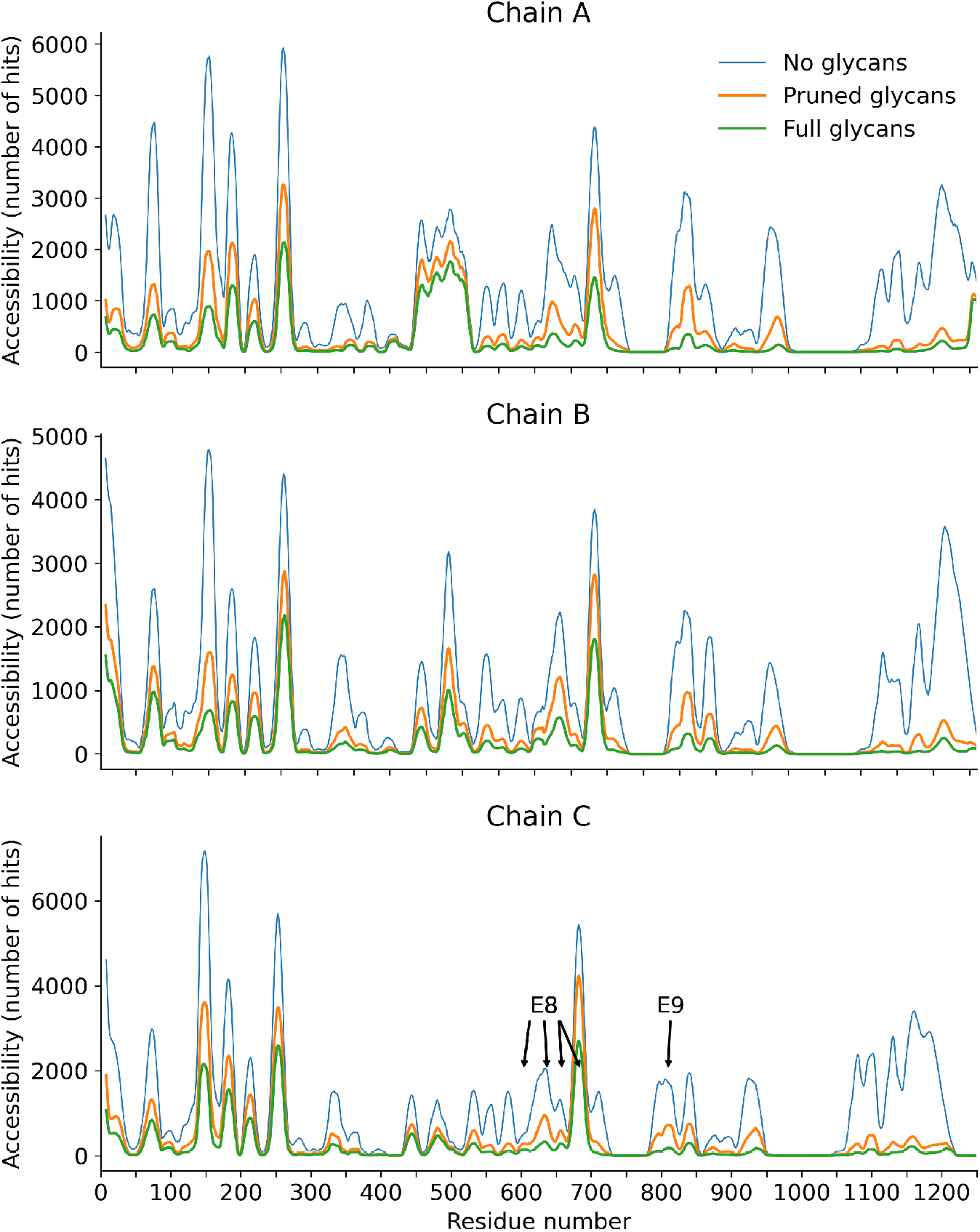
Impact of the glycosylation pattern on steric antibody accessibility. For each chain, the number of Monte Carlo rigid-body docking hits without glycans, with pruned glycans and with full glycans is shown. A rolling average over a 15 residue window was applied for legibility. Epitopes that undergo significant accessibility increases upon glycan pruning (E8 and E9) are indicated.

**FIG. S3.**
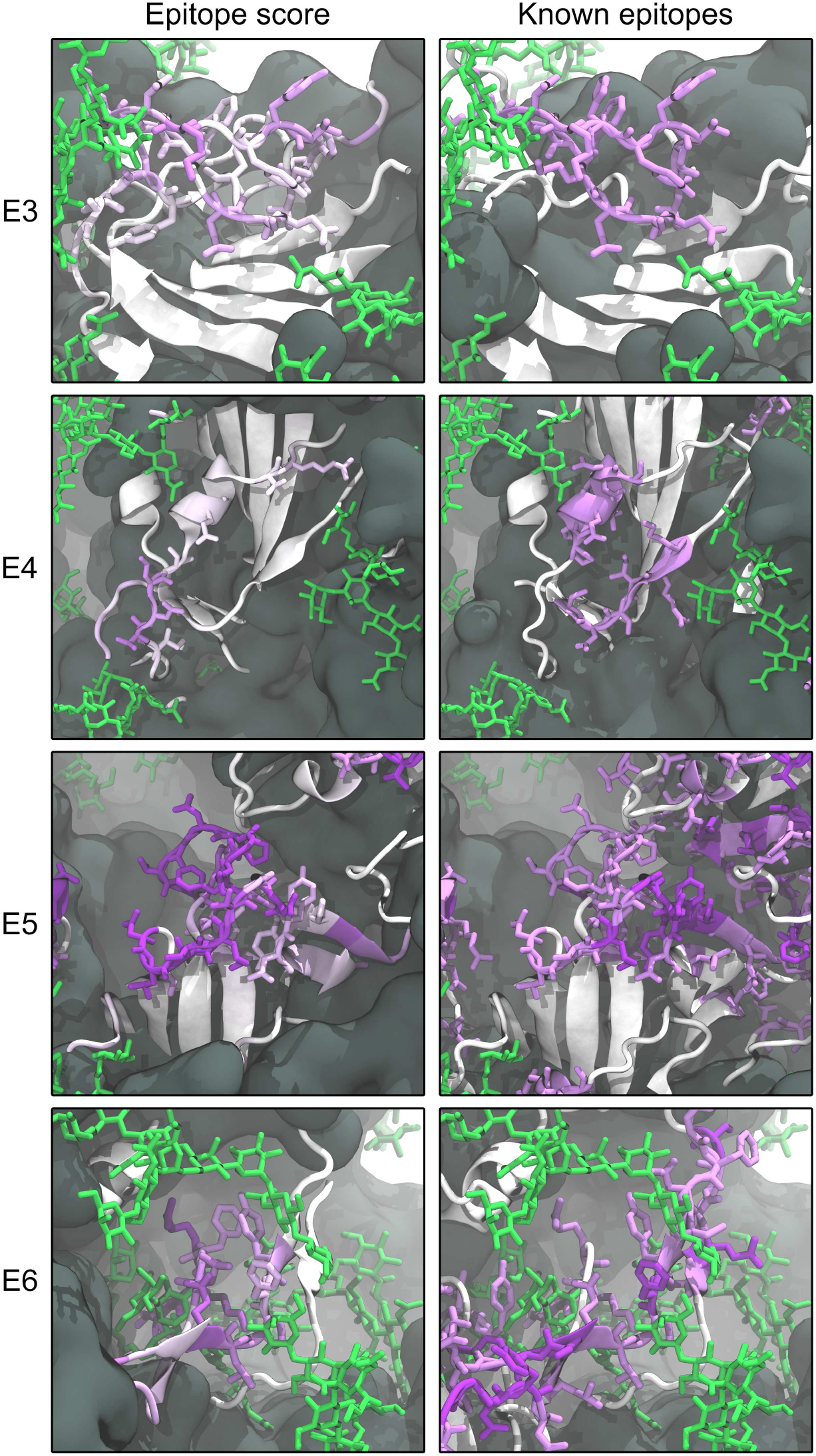
Comparison of the epitope candidates E3–E6 with previously characterized epitopes. Glycans are shown in green licorice representation. Left panels: Epitope candidates shown in cartoon representation with purple color intensity indicating epitope consensus scores. Residues with epitope consensus score >0.1 are shown in licorice representation. Right panels: Epitopes described in previous works shown in cartoon and licorice representation, with higher purple color intensity indicating reported binding to multiple distinct antibodies.

**FIG. S4.**
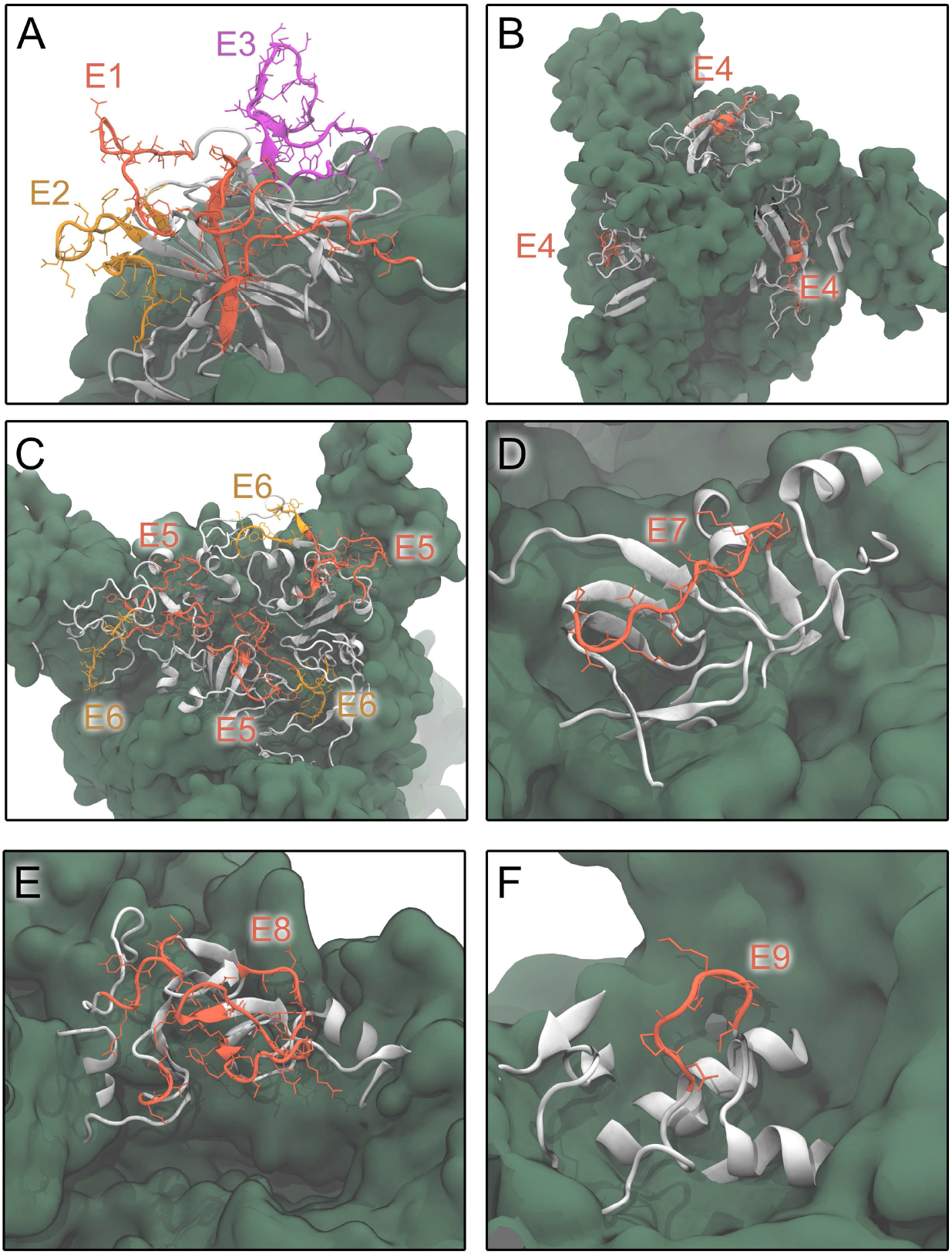
Location and structural features of the epitope candidates E1–E9 on the S surface. Epitope candidates are shown in red, orange and purple cartoon and licorice representation. Neighboring residues are shown in grey cartoon representation.

**FIG. S5.**
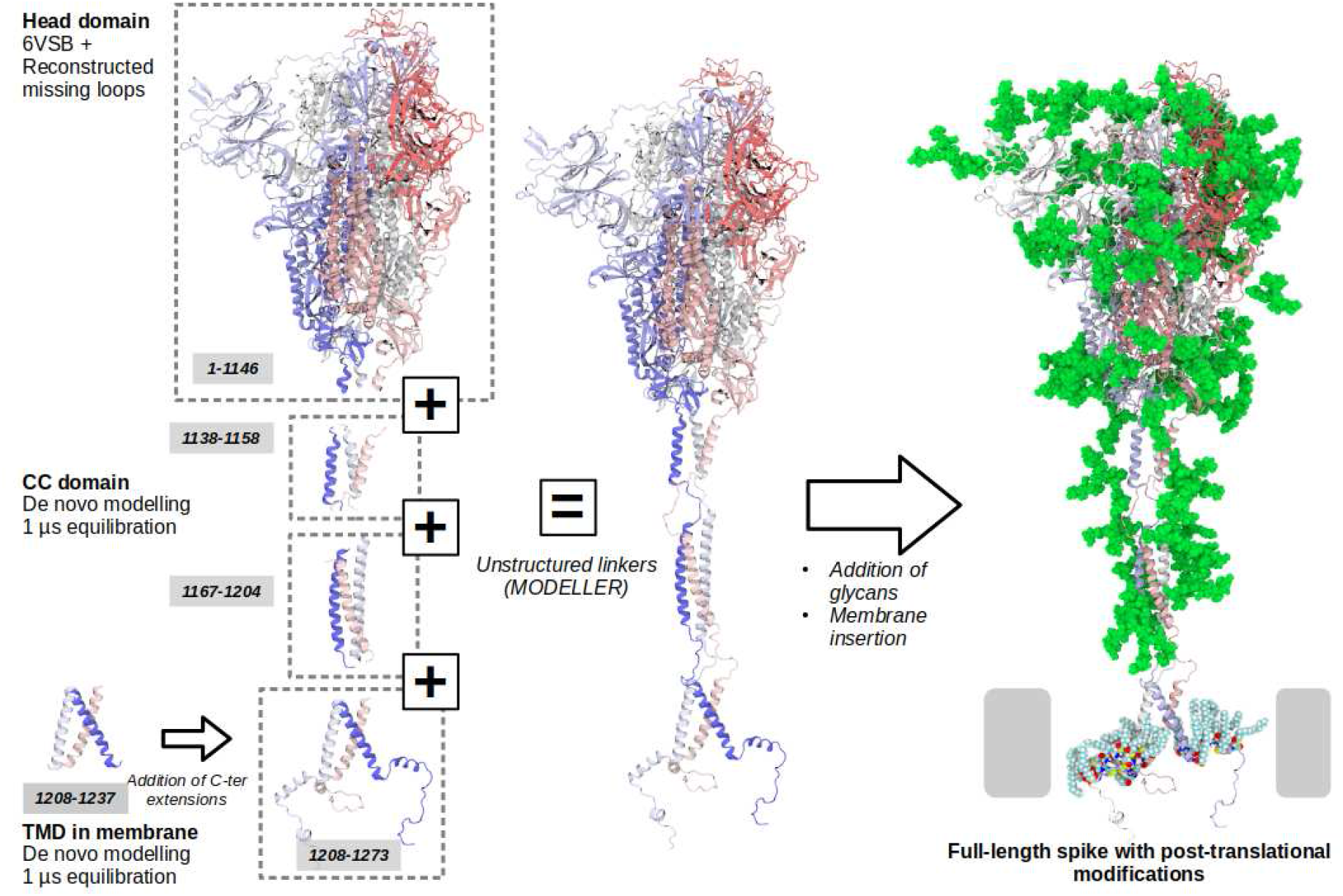
Schematic illustration of the strategy used to obtain an atomistic model of the full-length S protein. For clarity, we do not show the solvent and membrane.

**FIG. S6.**
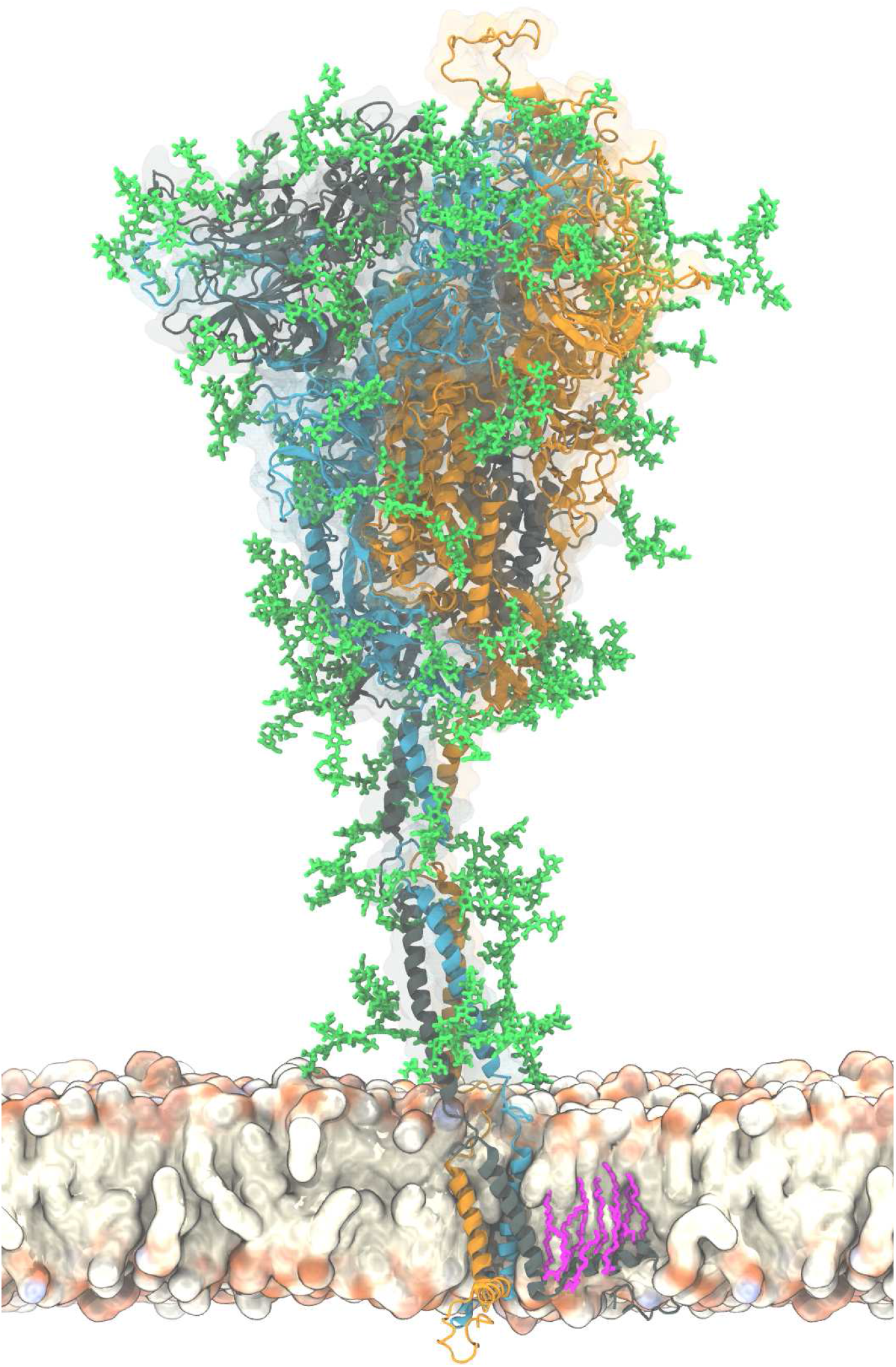
Atomistic model of the full-length membrane-embedded S protein shown in cartoon representation. The chains are differentiated by color. Palmitoylated cysteine residues are shown in pink licorice (only one chain shown for clarity). Glycans are shown in green licorice representation. We show a section of the membrane to highlight the transmembrane domain of S.

**FIG. S7.**
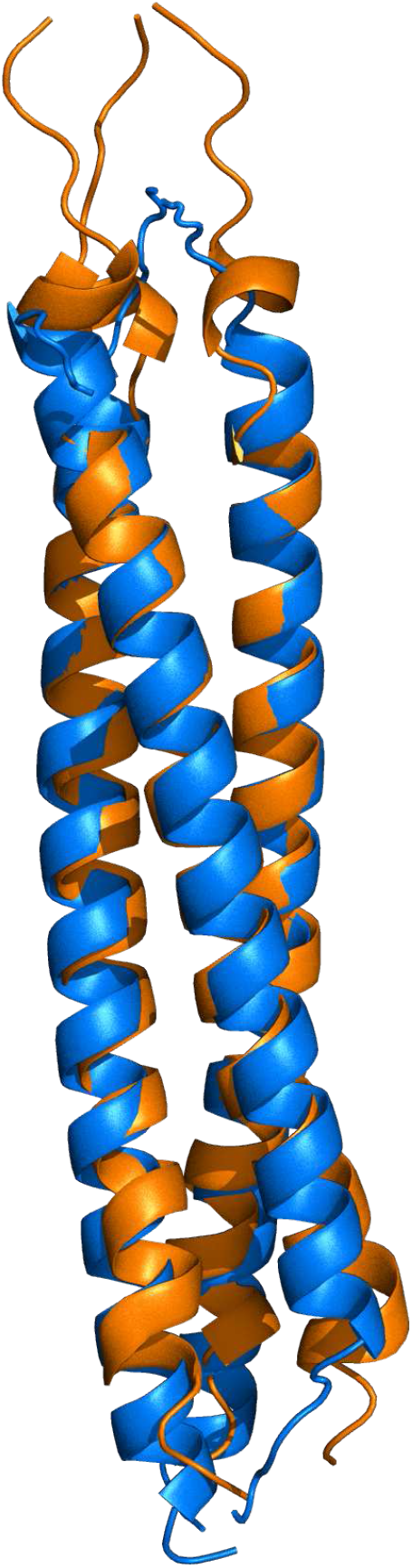
Rigid-body-aligned simulation structure of the HR2 coiled-coil (residues 1162-1214, blue) and SARS-CoV HR2 nuclear-magnetic-resonance solution structure 2FXP [31] (residues 1162-1212, orange).

**FIG. S8.**
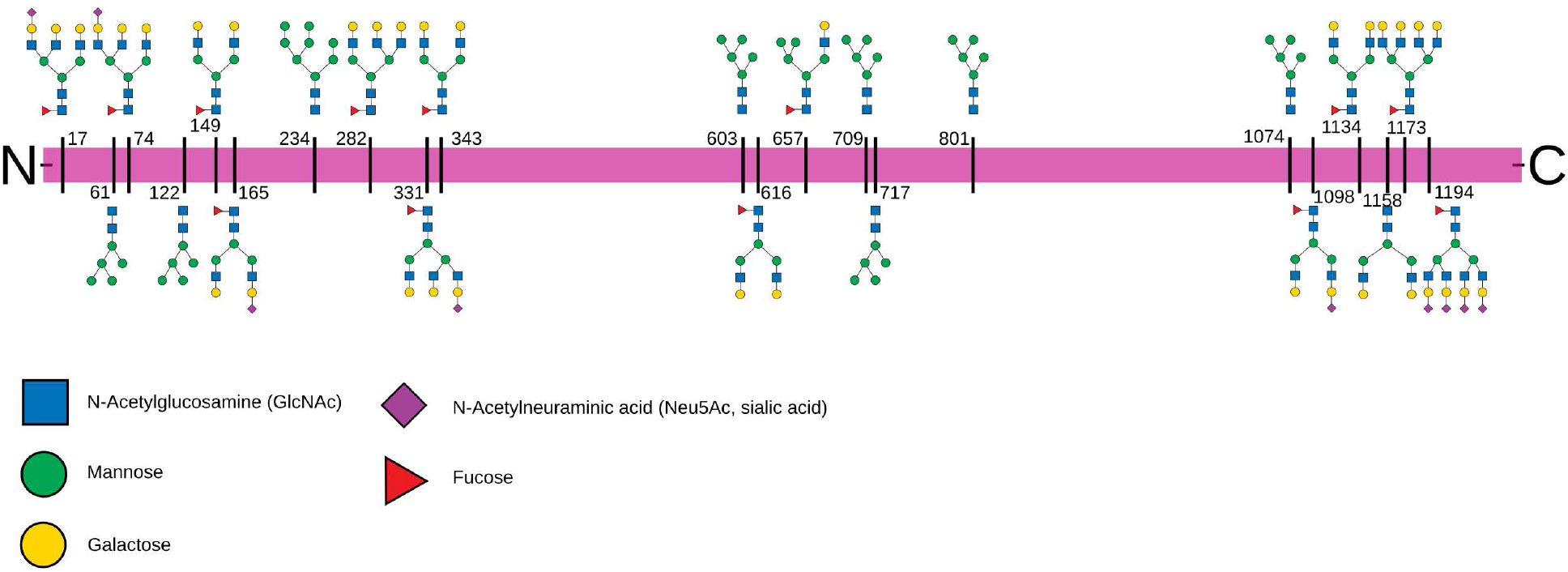
Glycosylation pattern of S. Sequons are indicated with the respective glycans in a schematic representation.

## SI MOVIE

Atomistic molecular dynamics simulation trajectory of four S proteins embedded in a membrane. The proteins and lipids are shown in surface representation. Glycans are represented by green van der Waals beads. Water and ions are omitted for clarity. 600 ns simulation time shown.

